# Pharmacological depletion of microglia prevents vascular cognitive impairment in Ang II-induced hypertension

**DOI:** 10.1101/2020.01.24.916650

**Authors:** Danielle Kerkhofs, Britt T. van Hagen, Irina V. Milanova, Kimberly J. Schell, Helma van Essen, Erwin Wijnands, Pieter Goossens, W. Matthijs Blankesteijn, Thomas Unger, Jos Prickaerts, Erik A. Biessen, Robert J. van Oostenbrugge, Sebastien Foulquier

## Abstract

**Rationale:** Hypertension is a major risk factor for cerebral small vessel disease, the most prevalent cause of vascular cognitive impairment. As we have shown, hypertension induced by a prolonged Ang II infusion is associated with increased permeability of the blood-brain barrier (BBB) and chronic activation of microglia. In this study we therefore aim to determine the contribution of microglia to hypertension-induced cognitive impairment in an experimental hypertension model by a pharmacological depletion approach.

**Methods:** For this study, adult *Cx3Cr1*^gfp/wt^x*Thy1*^yfp/0^ reporter mice were infused for 12 weeks with Angiotensin II or saline and subgroups were treated with PLX5622, a highly selective CSF-1R inhibitor. Systolic blood pressure (SBP) was measured via tail-cuff. Short- and long-term spatial memory were assessed during an Object Location task and a Morris Water Maze task (MWM). At the end of the study, microglia depletion efficacy and BBB leakages were assessed using flow cytometry and immunohistochemistry.

**Results:** SBP, heart weight and carotid pulsatility were increased by Ang II and were not affected by PLX5622. Short-term memory was significantly impaired in Ang II hypertensive mice, but not in Ang II mice treated with PLX5622. Histological and flow cytometry analyses revealed almost complete ablation of microglia upon CSF1R inhibition, while brain resident perivascular macrophages, were reduced by 60%. Number and size of BBB leakages were increased in Ang II hypertensive mice, but not altered by PLX5622 treatment.

**Conclusion:** Our results show that depletion of microglia, and less so PVMs, by CSF1R inhibition prevents short-term memory impairment in Ang II induced hypertensive mice. This novel finding supports the critical role of brain immune cells, and most in particular microglia, in the pathogenesis of hypertension-related cognitive impairment.

## Introduction

Cerebral small vessel disease (cSVD) is an age-related cerebral microangiopathy [1, 2]. It is expected that the prevalence of cSVD will increase in our aging society [2, 3]. cSVD is the leading cause of vascular cognitive impairment (VCI), an umbrella term that covers all cognitive disorders from mild cognitive impairment to vascular dementia [1, 4]. Hypertension is the major risk factor for the development of cSVD[2]. cSVD is associated with structural abnormalities on brain magnetic resonance imaging (MRI) including lacunes, white matter hyperintensities (WMH), cerebral microbleeds and enlarged perivascular spaces [5]. Despite a profound impact on human health, there is no specific treatment for cSVD [6, 7], mainly due to limited understanding of the disease’s pathobiology.

There is increasing evidence that blood-brain barrier (BBB) dysfunction plays a pivotal role in the pathophysiology of cSVD [8-12]. Under healthy conditions, the BBB functions in a well-regulated manner to ensure the provision of nutrients while protecting brain cells from blood constituents by forming a physical barrier [10, 13, 14]. The BBB is composed of vascular endothelial cells interconnected through tight junctions, flanked by pericytes, astrocytes, microglia cells and perivascular extracellular matrix. Proper interaction between its cellular and noncellular components is required to maintain a selective barrier function of the BBB [13, 14]. MRI studies have demonstrated higher BBB permeability in cSVD patients [8, 9] and this was associated with structural brain damage and cognitive impairment [8, 9, 15]. Pathological investigations have revealed the association of BBB leakages with WMH lesions and dementia [16, 17]. Leakage of plasma components into the parenchyma will elicit a local inflammatory response by Fc receptor-induced microglia activation, amongst others [18-20]. Microglia are brain resident myeloid cells which for their maintenance throughout the entire lifespan rely on their self-renewal capacity [21, 22], orchestrated by growth factor colony stimulating factor 1 (CSF1) and its receptor (CSF1R) [23]. In order to maintain physiological conditions, microglia are critical [24-27] by responding to conditions of tissue damage notably by clearing the accumulated debris [28, 29]. However, increased permeability of the BBB may lead to persistent microglia activation [30, 31], and potentially contribute to the progression of the pathology.

The use of Angiotensin II (Ang II) infusion has been previously associated with hypertension-induced cerebrovascular dysfunction including increased BBB permeability and neuroinflammation [32-34]. We have also shown earlier the presence of activated microglia in association with BBB leakages and short-term memory impairment in a prolonged Ang II infusion (12 weeks) model [35]. We hypothesize that the depletion of microglia protects against hypertension-induced cognitive dysfunction due to the absence of microglial activation. In this present study we aim to decipher the contribution of microglia to hypertension-induced cognitive impairment via their depletion using a highly selective CSF1R inhibitor.

## Material and Methods

### Animals

All animal experiments were approved by the regulatory authority of Maastricht University and were performed at Maastricht University in compliance with the national and European guidelines. *Cx3Cr1*^GFP/GFP^ mice (Jackson Lab 005582) were crossed with *Thy-1*^YFP/0^ mice (Jackson Lab 003782) to generate *Cx3Cr1*^GFP/WT^ x *Thy1*^YFP/0^ mice (abbreviated Tg mice) for microglial and neuronal visualization. Animals were kept on a normal 12h day-night cycle. All mice were allowed access *ad libitum* to water and mouse chow. At the start of the experiment, 3-months old male Tg mice were fed with either PLX5622 laced chow (Plexxikon; CSF1R tyrosine kinase inhibitor; 1200 ppm) or control chow for 12 weeks. At the same time, all mice were equipped with osmotic minipumps (Alzet model 2006, Direct Corp., Cupertino, CA, USA) for the delivery of Ang II (1 μg/kg/min subcutaneously) or saline. Osmotic minipumps were implanted *s.c.* under isoflurane anesthesia and were replaced after 6 weeks with new minipumps for a total delivery time of 12 weeks. In total, 45 mice were included in the 4 groups (11-12 per group, Figure 1A). Three animals died or were sacrificed prior to the end of the study due to haemorrhage in the thoracic cage (Ang II, vehicle), peritonitis due to a missed injection (Ang II, PLX5622) and severe weight loss after blood pressure measurement (Saline, PLX5622). Mice used for histology received an *i.v.* injection of 70kDa-dextran Texas Red (100 μL; 2,5 mg/mL in sterile NaCl 0,9%, D1864, Thermo Fisher), while under isoflurane anesthesia, to detect the presence of BBB leaks by histology. The injected dextran was allowed to circulate for 3 min before mice were euthanized by exsanguination. A second series of mice was euthanized after blood collection and intra-cardiac perfusion with PBS and heparin for FACS analysis of blood and brain samples.

**Fig. 1:**
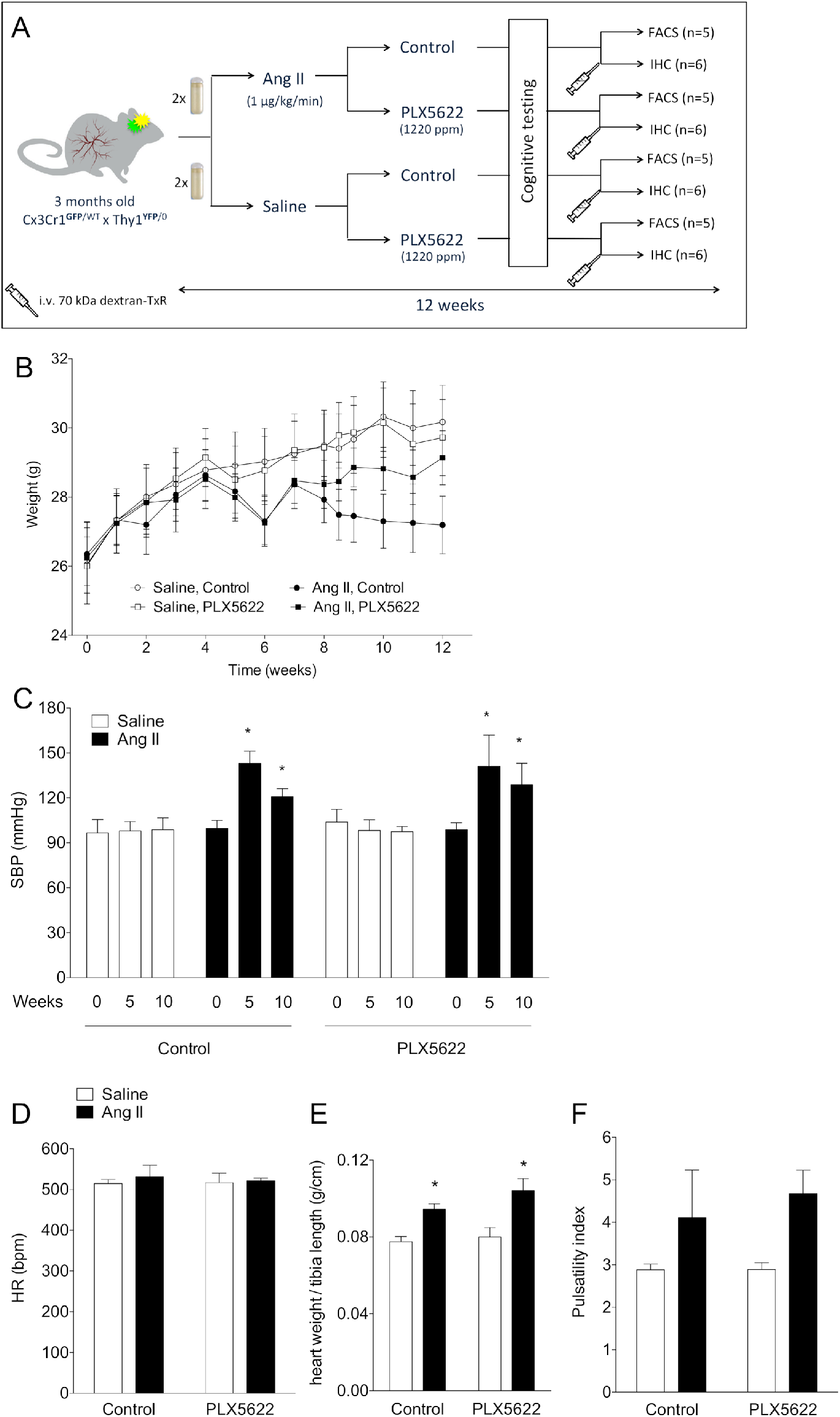
Study design, body weight and cardiovascular parameters. Design of the study **(A);** Body weight progression over the study period (2-W ANOVA p_int_ >0.05; p_time_< 0.001; p_groups_ < 0.001) **(B);** Systolic blood pressure values at baseline (week 0), mid-term period (week 5) and final-term period (week 10) (2-W ANOVA p_int_ < 0.001; p_time_< 0.001; p_AngII_ < 0.05; Tukey’s multiple comparison test: *p < 0.05 vs. Saline) **(C);** Heart rate at week 12 (2-W ANOVA p_int_ > 0.05; p_plx5622_ > 0.05; p_AngII_ >0.05) **(D);** Cardiac hypertrophy (heart weight/tibia length) (2-W ANOVA pint > 0.05; p_PLX5622_ > 0.05; p_AngII_ < 0.001; Tukey’s multiple comparison test: *p < 0.05 vs. Saline) **(E);** Carotid pulsatility index at week 12 (2-W ANOVA p_int_ > 0.05; p_plx5622_ > 0.05; p_AngII_ = 0.01)**(F)**. Ang II (full boxes/bars), control (empty boxes/bars). n = 9–11 per group. (FACS: Fluorescence Assisted Cell Sorting; IHC: immunohistochemistry)

### Cardiovascular phenotyping

Systolic BP was monitored before minipump implantation and after 5 and 10 weeks in awake mice using tail-cuff plethysmography (CODA, Kent Scientific) as previously described [35]. Before sacrifice, ECG and carotid blood flow velocity signals were acquired non-invasively from the anaesthetized mice using ultrasound flow velocity Doppler (20 MHz probe, DFVS, Indus Instruments, Webster, TX, USA). Mice were laid down in supine position on a temperature-controlled ECG board (Rodent surgical monitor; Indus Instruments, Webster, TX, USA). Body temperature was monitored with a rectal probe and maintained at 37°C. Three blood flow velocity records were saved per carotid. Heart rate (HR, bpm) and blood flow signals were analysed offline using the Indus Instruments Doppler Signal Processing Workstation and were averaged per mouse. Carotid pulsatility index (PI) was calculated as PI = [Peak systolic velocity – End diastolic velocity]/Mean velocity.

### Cognitive tests

Long-term spatial memory was tested in the Morris Water Maze (MWM) [36]. Shortly, the swim tank was divided into four quadrants and a platform was located in a fixed location. A video camera automatically recorded the mice movements (via a tracking system EthoVision, Noldus). Mice were subjected to the following testing schedule for MWM: spatial navigation with acquisition (days 1-4) and a probe trial (day 5). In all testing procedures the mice were given four trials per day with an inter-trial interval of 10 minutes and using four different starting positions. Each trial started with the mouse facing the wall of the pool and ended when the mouse reached the platform, or after 60 s. At day 5, a probe trial was conducted in which the platform was removed from the tank and the mouse was allowed to swim in the tank for 60 s. The swimming distance to reach the platform (cm) during test trials as well as the swimming speed and the time spent in the target quadrant during the probe trial (seconds), were obtained by Ethovision.

Short- and long-term memories were assessed in the two weeks preceding sacrifice. Short-term spatial memory was tested using an object location task (OLT) at 1 h intertrial intervals[35]. Briefly, this 2-trial task consisted of a learning (T1) trial and a test trial (T2). In T1, a set of two identical objects were placed symmetrically in the middle of a circular arena, which the mouse was allowed to explore freely for 4min. After a 1 h interval spent in their home cage, the mice were placed in the arena for T2; in this 4 min-trial, one of the objects (right or left) was moved to a different location (front or back), while all other stimuli were kept the same. Mice will spend more time exploring the moved object than the stationary object if they remember the previous location. The time spent exploring each object was scored manually on a computer by an experimenter blind to the experimental groups. Trials were excluded from the analysis when the total exploration time was inferior to 6 seconds. The discrimination index d2 assesses whether the mouse spends more time at the novel location than at the familiar location (the difference between location exploration times divided by the total exploration time). Functional spatial short-term memory is reflected by a d2 index higher than zero (both objects equally explored) [37].

### Blood and brain FACS

Heparinized whole blood was used for flow cytometry using the following antibody cocktail: CD45-PerCP (Biolegend 103130), CD3-eFLUO450 (eBioscience 48-0032), NK1.1-PE (BD557391), Ly6G-APC-CY7 (BD560600), CD11b PE-CY7 (BD552850), Ly6C-APC (Miltenyi 130-093-136), CD19-APC-H7 (eBioscience 47-0193) and Siglec-F-PE (BD552126). Further, cells from the whole brain were isolated for FACS. Anesthetized mice were perfused with PBS to remove the peripheral blood. Subsequently, brains were dissected, mechanically dissociated and digested with a collagenase mix (including collagenase IX, collagenase I, DNAse and RPMI-HEPES). Immune cells were separated using a Percoll-gradient and stained with CD45 PerCP (Biolegend 103130), CD11c PE-Cy7 (eBioscience 25-0114)), F4/80 (Biolegend 123116), CD11b BV510 (Biolegend 101245), CD3 BV421 (eBioscience 48-0032-82), CD19 BV421 (Biolegend 115538), Ly6G BV421 (eBioscience 48-3172). All samples were measured with a FACS-Canto II (BD Biosciences). Results were analysed with FACSdiva version 8 (BD Biosciences).

### Immunohistochemistry

Brains were post-fixed with 4% paraformaldehyde overnight at 4°C, washed with Tris Buffered Saline (TBS) and stored in TBS containing 0.1% sodium azide at 4°C. Coronal sections (thickness 50 μm) were prepared using a vibratome (VT1200S, Leica). Free-floating sections were thoroughly washed with 0.3% Triton-TBS followed by antigen retrieval with target retrieval solution citrate pH6 (1:10, Dako S2031) for 20 minutes at 80°C. After blocking with 1% of bovine serum albumin in 0.1% Triton-TBS, sections were incubated with primary antibodies anti-Iba1 (rabbit polyclonal, 1:1000; Wako 019-19741), anti-CD206 (MR5D3, rat polyclonal, 1:200, Bio-Rad MCA2235), anti-MBP (Myelin Basic Protein, rat polyclonal, 1:1000 Millipore MAB 386) or antimouse IgG (Biotin-SP AffiniPure, donkey polyclonal, 1:100, Jackson ImmunoResearch N715-065-150) overnight at 4°C. After 3 washes, the sections were incubated with secondary antibodies including donkey-anti-rat biotin (1:400, Jackson ImmunoResearch 712-065-150) or donkey-anti rabbit biotin (1:400, Jackson ImmunoResearch 711-065-151), followed by conjugation with Streptavidin-AF647 (1:500, ThermoFischer S32357). Brain sections were mounted on gelatin-coated microscopic slides using fluorescence mounting medium (Dako S3023) and examined with a slide scanning microscope (Nikon Eclipse Ti-E) and a confocal microscope (Leica SPE).

### Quantifications of microglia, perivascular macrophages, blood brain barrier leakages, myelin integrity and neuronal loss

All analyses were performed by a blinded investigator using ImageJ (Fiji Distribution, NIH). Microglia were counted as Cx3Cr1^+^/Iba1^+^ cells on 6 randomly selected volumes within the neocortex (x=366; y=366; z=20 μm; n=5-6). Perivascular macrophages (PVMs) were counted as CD206^+^ cells[38] on 8 volumes selected to include cortical penetrating vessels within the neocortex (x=275; y= 275; z=22 μm; n=5-6). BBB permeability was assessed by determining the level of extravasation of the 70kDa-dextran probe and plasmatic IgG proteins into the brain parenchyma as shown previously [35]. Identification of IgG leakages was performed morphometrically by one investigator, who was blinded to the experimental groups. BBB leakages were defined as a signal with an intense core and diffuse borders. A series of brain slices (6 slices per brain; n=6 mice) were screened to identify and localize IgG and dextran leakages (Nikon Ti-Eclipse slide scanner). Z-stack images of all identified leakages were then acquired by confocal microscopy (x= 175; y= 175; z=20 μm). The fluorescent signal of each leakage was then analyzed in ImageJ to quantify the leakage size in mm^2^. MBP signal in the corpus callosum, a myelin-rich area, was used to assess changes in myelin composition in the different experimental groups. Corpus callosum thickness and MBP signal intensity were measured at 3 defined locations in the lateral and medial corpus callosum (bregma level: +0.7 mm, 2 slices per brain, n=6). The intensity of the myelin signal was examined in 8 leakages versus contralateral brain regions without leakages. Image stacks (x=550; y=550; z=30 μm) of leakages were acquired by confocal microscopy.

### Chemicals

All chemicals were from Sigma-Aldrich (Zwijndrecht, The Netherlands) unless otherwise specified. PLX5622 was provided by Plexxikon Inc. and formulated in standard chow (19% protein, 5% fat, 5% fiber, 0.2%).

### Statistical analyses

All statistical analyses were performed with GraphPad Prism 6 software. Data are expressed as standard error of the mean (SEM). The normality distribution was tested with the Shapiro-Wilk test. Two-way ANOVA tests were performed with Ang II and PLX5622 treatments as independent variables, followed by Tukey’s or Sidak multiple comparison post-hoc tests between individual groups. A p value < 0.05 was considered as statistically significant.

## Results

### CSF1R inhibition and cardiovascular phenotype

PLX5622 treatment had no impact on body weight progression. Ang II infusion decreased the body weight in both control and PLX5622 groups compared to Saline infused mice (27,2±0.8 g and 29.1±0.8 g in Ang II groups vs 30.2±1.1 g and 29.7±1.1 g in Saline groups respectively at week 12; Fig. 1B, group effect p_Ang II_ < 0.05). Systolic blood pressure was increased by Ang II at 5 weeks in both PLX5622 and vehicle treated groups and remained elevated at 10 weeks in both groups (Fig. 1C, p_Ang II_ <0.001). Heart weights, normalized to tibia lengths, as well as carotid pulsatility indices were significantly increased in the Ang II compared to Saline groups and not changed by PLX5622 treatment (Fig. 1E, p_AngII_ <0.001 and Fig. 1F, p_AngII_ <0.05, respectively).

### CSF1R inhibition and number of microglia, perivascular macrophage number and circulating immune cells

Flow cytometry analysis of CD45^int^Cx3Cr1^hi^CD11b^hi^ cells revealed effective depletion of microglia in whole brains by the CSF1R antagonist PLX5622 (−94% in Saline groups; −83% in Ang II groups; p_PLX5622_<0.0001) (Fig. 2A). This was confirmed by immunohistochemistry showing profoundly decreased density of Cx3Cr1^+^Iba-1^+^ cells in the cerebral cortex of PLX5622 treated mice (−99% in Saline groups; −92% in Ang II groups; p_PLX5622_<0.001) (Fig. 2B). CD206^+^ PVMs numbers, surrounding cortical penetrating arterioles, were reduced as well in the PLX5622 treated groups (67% in Saline groups; −56% in Ang II groups; p_PLX5622_<0.001) (Fig. 2C). Treatment with PLX5622 did not influence CD45^+^ cells, CD19^+^ B-cells and CD3^+^ T-cells in the blood (Fig. 3A-C). CD45^+^Cx3Cr1^+^Ly6C^low^ non-classical monocytes were reduced after 12 weeks of PLX5622 treatment (−64% in Saline group; −69% in Ang II group; p_PLX5622_<0.001), whereas the number of CD45^+^Cx3Cr1^+^Ly6C^high^ classical monocytes did not change (Fig. 3D-F).

**Fig. 2:**
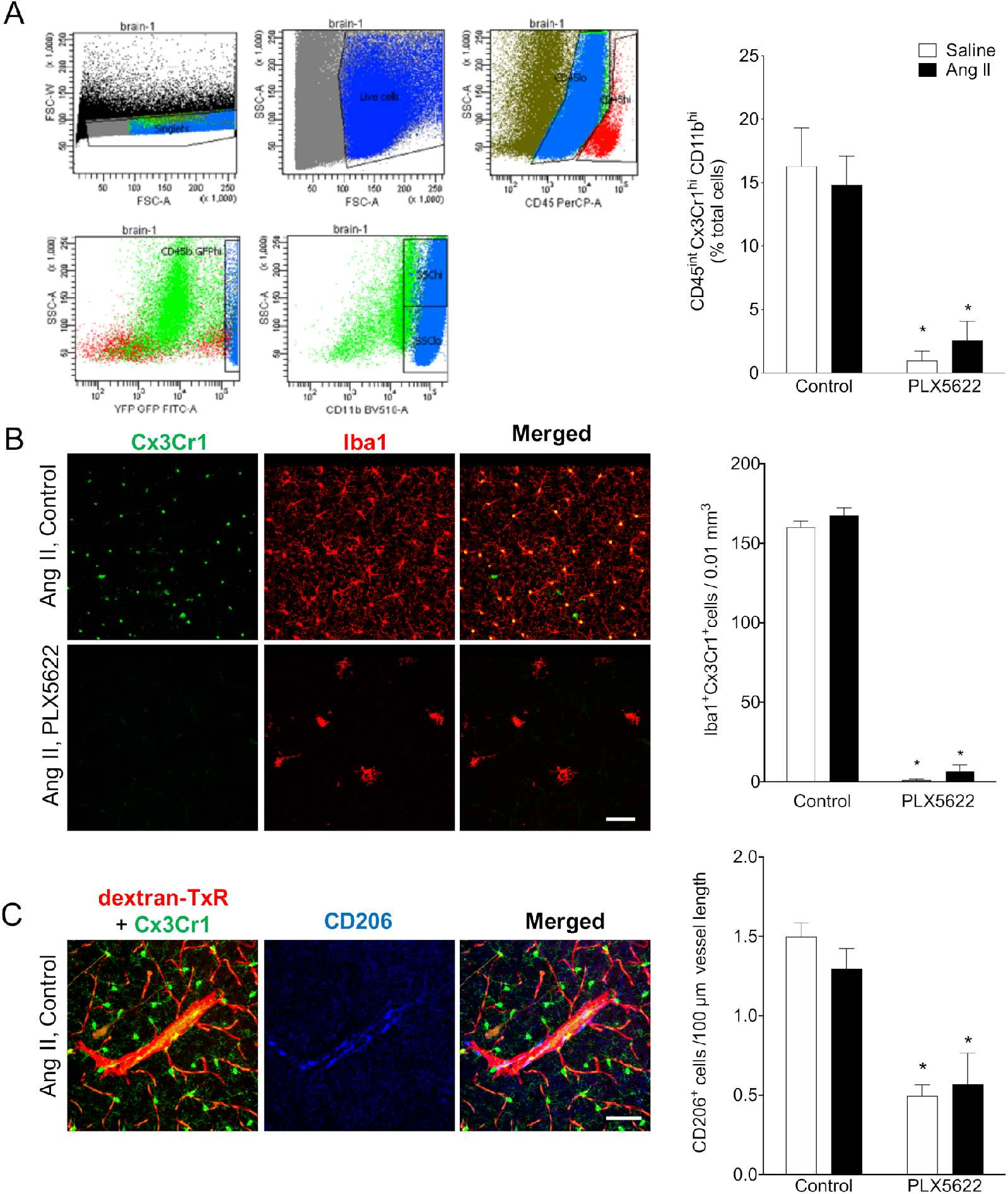
Impact of CSF1R inhibition on microglia and perivascular macrophage densities. Representative flow cytometry gating of microglia population (CD45^int^,Cx3Cr1^hi^,CD11b^hi^) **(A, left figure)** and bar graph of microglia cells as percentage of total cells (2-W ANOVA p_int_ >0.05; p_plx5622_ < 0.001; p_AngII_ > 0.05; Tukey’s multiple comparison test: *p < 0.05 vs. control) **(A, right figure);** representative pictures of Cx3Cr1-positve cells (green) and Iba-1 postive cells (red) in cortical areas of Ang II-infused control **(B, upper row)**; and Ang II-infused PLX5622 **(B, lower row)** (20x magnification, scale bar = 50 μm); microglia densities in cerebral cortex (2-W ANOVA p_int_ >0.05; p_plx5622_ < 0.001; p_AngII_ = 0.02; Tukey’s multiple comparison test: *p < 0.05 vs. Control) **(B, right);** representative picture of CD206^+^ staining in cortical areas of Ang II-control mice **(C);** Left column: of Cx3CR1^+^ cells (green) and dextran- TxR (red). Middle column CD206^+^ elongated cells along the wall of penetrating neocortical vessels (blue). Right column: composite image. (40x magnification, scale bar = 50 μm). Perivascular macrophage densities in cerebral cortex (2-W ANOVA p_int_ >0.05; p_plx5622_ < 0.001; p_AngII_ > 0.05; Tukey’s multiple comparison test: *p < 0.05 vs. Control) **(C, graph**). n = 9–11 per group

**Fig. 3:**
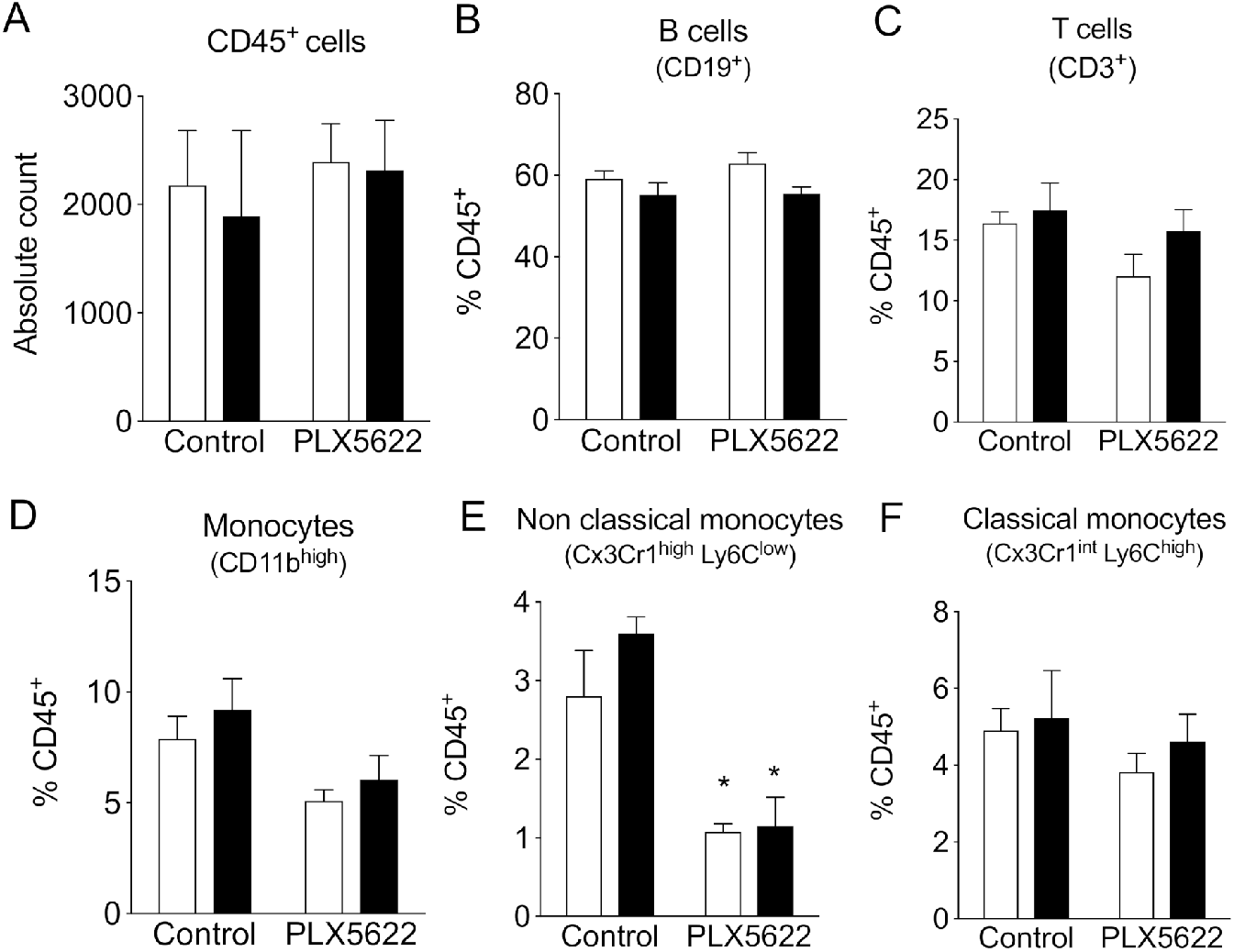
Impact of CSF1R inhibition on circulating immune cells. Cell counts of circulating CD45^+^ cells (A), CD19^+^ B-cells (B), CD3^+^ T-cells (C), monocytes (D), of the Ly6C^low^ (E) and Ly6C^high^ subset (F), measured with flow cytometry. 2-way ANOVA; Tukey’s multiple comparison post-test (*: p < 0.05 vs Vehicle). Ang II (full boxes/bars), control (empty boxes/bars). n = 3–4 per group.

### CSF1R inhibition attenuates short-term memory impairment in hypertensive mice

Mice were first tested on the Morris water maze to evaluate learning and spatial memory[39]. There was no overall effect of PLX5622 treatment and/or Ang II infusion on the escape distance during the training trials. The distance to reach the platform was however increased for the saline PLX5622 group vs Saline controle group but only at day 2. (Fig. 4A). For the probe trial, swimming speed was not different between the groups, which is indicative of normal motor behaviour. Both Ang II and PLX5622 did not influence the time spent in the target quadrant (Fig. 4B, C). Thus, long-term memory performance during the MWM task was not affected by Ang II and/or PLX5622 treatment.

**Fig. 4:**
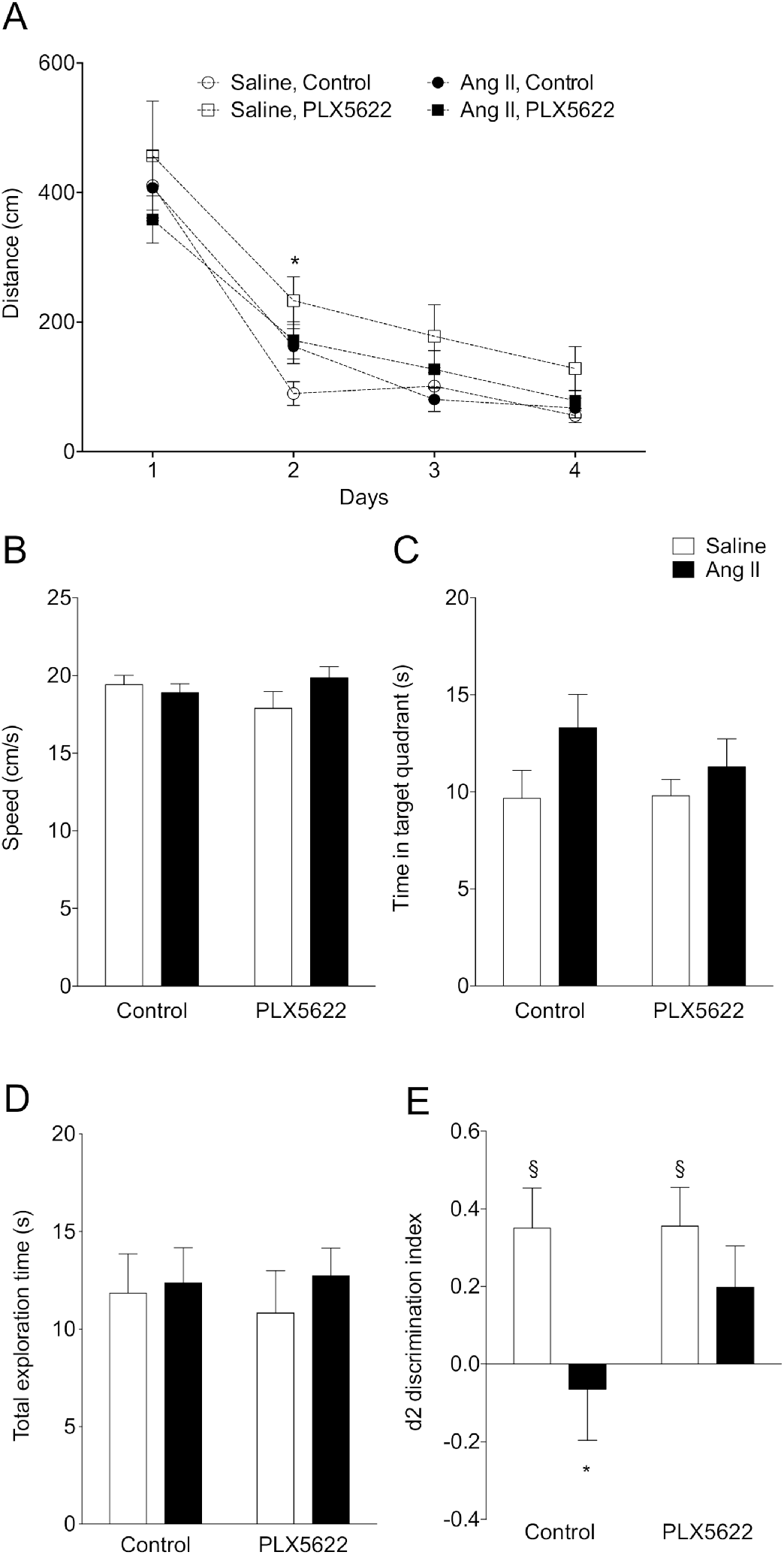
Cognitive performance. Long-term memory was assessed using the Morris Water Maze Task (A,B,C). Swimming distance to reach the platform (2-W ANOVA p_int_ >0.05; p_time_< 0.001; p_AngII_ < 0.05; Tukey’s multiple comparison test: *p < 0.05 vs. Control) (A); swimming speed (2-W ANOVA pint > 0.05; p_plx5622_ > 0.05; p_AngII_ >0.05) (B); time spent in target quadrant during the probe trial (2-W ANOVA p_int_ > 0.05; p_plx5622_ > 0.05; p_AngII_ >0.05) (C). Short-term memory assessed using an Object Location Task at 12 weeks (D,E). Total exploration time (2-W ANOVA p_int_ > 0.05; p_plx5622_ > 0.05; p_AngII_ >0.05) (D); and discrimination index d2 (2-W ANOVA pint > 0.05; p_PLX5622_ > 0.05; p_AngII_ < 0.05; Sidak’s multiple comparison test: *p < 0.05 vs. Saline Control; Two tailed t-test: §: p <0.05 vs d2=0) (E). n = 9–11 per group.

In the object location task, the total exploration times did not differ between groups, indicating again that motor behavior is normal in all animals (Fig. 4D). Normotensive mice spent more time exploring the object at the novel location as indicated by the positive discrimination index d2 (Saline, Control: p=0.01; Saline, PLX5622: p=0.02). PLX5622 did not affect task performance in normotensive mice. Vehicle treated Ang II-infused mice were unable to discriminate between the two objects with d2 different from the Saline group and not different from zero, i.e. chance performance (p=0.69). However, Ang II-infused mice treated with PLX5622 performed significantly better compared to untreated Ang II-infused mice, as d2 did not differ from its respective Saline group. The discrimination index d2 was however not completely back to normal as it was still not statistically different from 0. (Fig. 4E).

### CSF1R inhibition and Ang-II induced blood-brain barrier leakage and myelin integrity

BBB permeability was assessed by the extravasation into the brain parenchyma of the 70kDa-dextran probe and plasmatic IgG proteins (Fig. 5A). Both the total number (p_AngII_ = 0.001; Fig. 5B) and average size of BBB leaks, as judged from 70 kDa dextran and IgG extravasation (p_AngII_ = 0.02 and p_AngII_ = 0.003 respectively; Fig 5C,D) were significantly increased in the Ang II groups. Treatment with PLX5622 did not influence number or size of BBB leaks. The local decrease in microglia density at the leakage site in the PLX5622 treated groups was equivalent to the observed global depletion (p_plx5622_ < 0.001) and there was no effect of Ang II on the microglia density at the leakage sites (Fig. 5E). Myelin intensity in leakage sites in the PLX5622 treated animals was not significantly different between the saline and Ang II groups (p=0.7) (Fig 6A,B). The size of the corpus callosum (Fig. 6C) and the intensity (data not shown) of the myelin signal in corpus callosum was not different between the study groups.

**Fig. 5:**
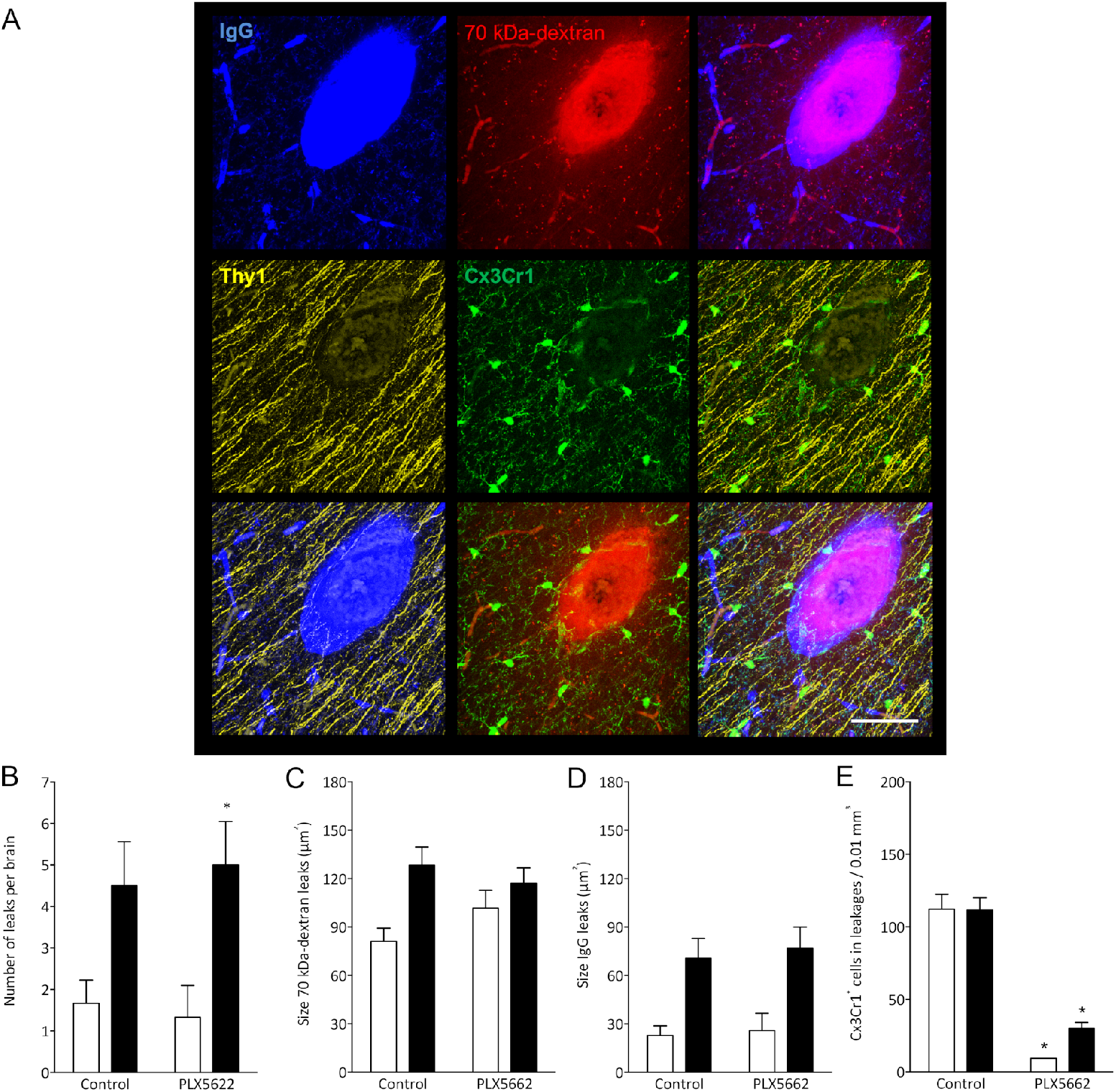
Number and size of blood-brain barrier leakages. Representative picture of a leakage identified by 70 kDa dextran-Texas Red and IgG-positive stainings (blue), Thy-1^+^ axons (yellow) and Cx3Cr1^+^ cells (green) (63x magnification scale bar = 50 μm) (A); Number of leakages per brain identiefied in 6 brain sections (2-W ANOVA pint > 0.05; p_PLX5622_ > 0.05; p_AngII_ = 0.001; Sidak’s multiple comparison test: *p < 0.05 vs. Saline) (B); Average size of the 70 kDa dextran leakages (2-W ANOVA p_int_ > 0.05; p_plx5622_ > 0.05; p_AngII_ = 0.02) (C); Average size based of IgG^+^ leakages (2-W ANOVA p_int_ > 0.05; p_plx5622_ > 0.05; p_AngII_ = 0.003) (D); Density of Cx3Cr1^+^ cells in leakages (2-W ANOVA p_int_ >0.05; p_plx5622_ < 0.001; p_AngII_ = 0.63; Sidak’s multiple comparison test: *p < 0.05 vs. Control) (e); in all experimental groups. n = 9–11 per group.

**Fig. 6:**
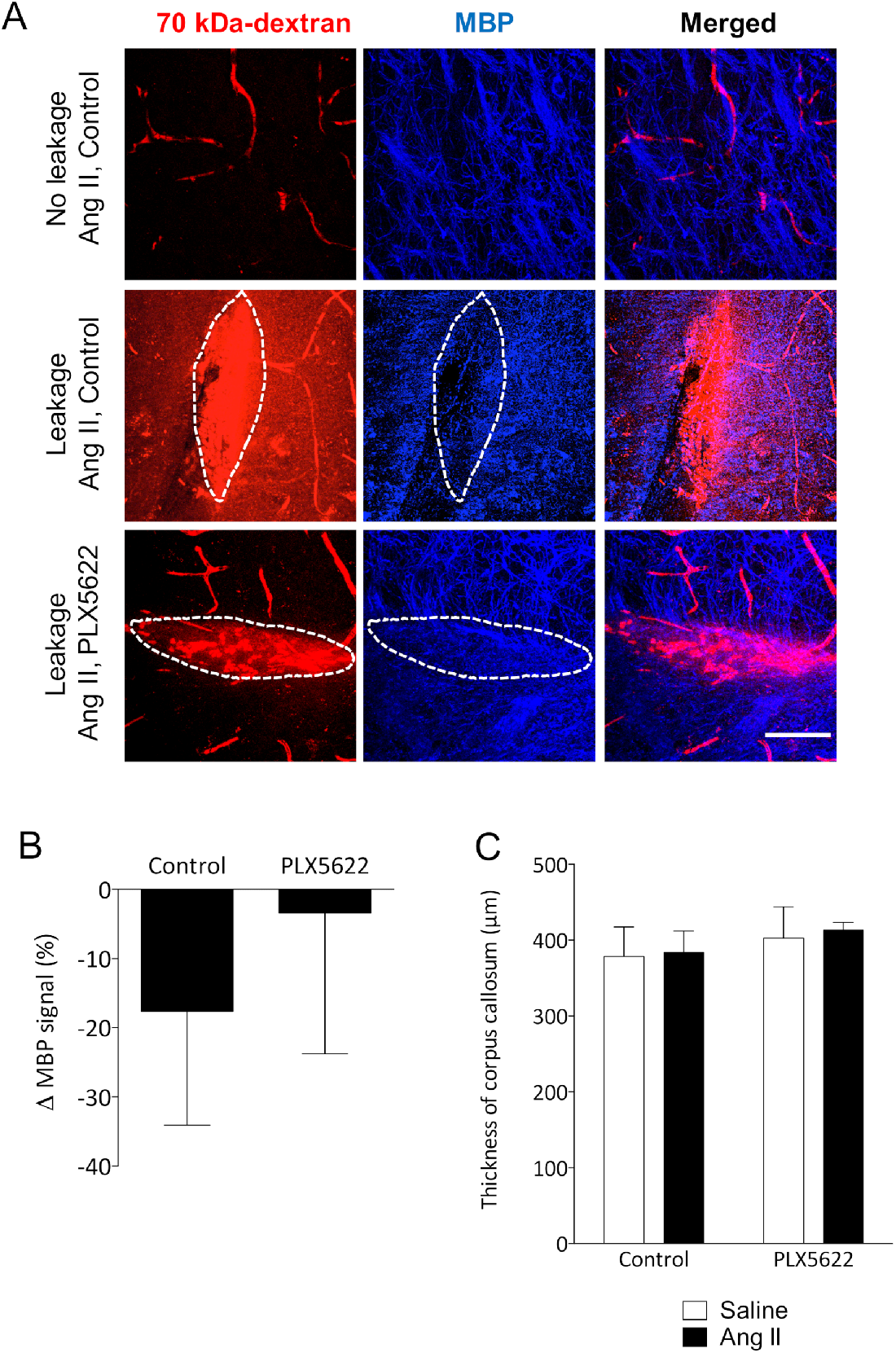
Myelin loss at leakage sites. Myelin Basic Protein (MBP) signal (blue) in the absence (upper row) and presence (middle and botom row) of 70 kDa dextran leakages (red) in Ang II Control (middle row) and PLX5622 treated Ang II mice (bottom row) (63x magnification, scale bar = 50 μm) **(A)**; Change of MBP signal in Dextran-positive vs Dextran-negative areas in Ang II mice treated with PLX5622 compared to Control; n=4 leakages per group **(B)**; Thickness of medial corpus callosum **(C)**.

## Discussion

An impaired BBB permeability induced by chronic hypertension is associated with microglia activation and short-term memory impairment [32-35]. In the present study we aimed to determine the contribution of microglia cells to hypertension-induced cognitive impairment using a pharmacological depletion approach. Our main finding is that short-term memory impairment caused by Ang II-induced hypertension was absent after microglia / PVM depletion, in support of a critical role of microglia in the pathogenesis of VCI.

To determine the effects of microglia depletion on cognitive function, short- and long-term spatial memory were assessed in an OLT and a MWM, respectively. In the OLT, we found that hypertensive mice had impaired short-term memory, as they could not discriminate between the new and the old object location, confirming earlier work by us [35], and others, based on similar tasks [33]. Treatment with the CSF1R inhibitor PLX5622 partly obviated the hypertension associated impairment of short-term memory without altering the normal cognitive function of control mice. In the Morris Water Maze task however, both groups of Ang-II infused mice showed a normal spatial learning and memory as observed previously [40]. The overall performance of the animals in the PLX5622 groups was also not altered, although post-hoc testing revealed an increased distance to reach the platform in the Saline PLX5622 group at day 2. It is debatable whether this statistical significant effect has any biological significance taking into account the overall lack of effect of PLX5622 on normal behavior [41].

The observed attenuated hypertension-induced short-term memory impairment mediated by CSF-1R inhibition is a novel finding. This results aligns with findings in other brain disease models as microglia depletion by CSF1R inhibition has proven to be effective to prevent radiation induced short-term memory impairment in mice [42] and to improve the cognitive function in a mouse model of Alzheimer’s disease [41]. In analogy to previous short- and long-term CSF1R inhibition studies, the overall behaviour and cognitive function of control mice were unaltered [23, 41-43]. Furthermore, dampening microglial reactivity using minocycline, was benefical for subcortical white matter functioning and cognitive performance in an animal model of chronic cerebral hypoperfusion[44, 45], strengthening the importance of targeting microglia to limit the impact of cerebrovascular diseases.

In accordance with our prior study [35], we found an increased BBB permeability in Ang II infused hypertensive mice, which was not altered by PLX5622 (Fig. 5). Increased BBB permeability causes leakage of plasma components which can induce microglia activation by binding the Fc receptors on these brain resident immune cells [18, 33, 46]. IgG leakage in the brain parenchyma is associated with an exacerbated neuroinflammatory response as shown by the increased density and soma size of microglia in different brain regions [35]. Quantification of the microglia density around the leakage sites revealed overt depletion of microglia cells surrounding the leakage in the PLX5622 treatment groups, demonstrating that PLX5622 was also effective at the site of leakages. This suggests that the expected BBB leakage-associated neuroinflammatory response is dampened by the treatment.

Emerging evidence from animal models and clinical studies on cSVD points to a clear association between increased blood brain barrier permeability and cognitive impairment [11, 15, 30]. Our results confirm this association between BBB leakages and cognitive impairment in the untreated hypertensive group and moreover adds new knowledge by demonstrating that CSF1R targeted depletion of microglia can prevent cognitive decline in hypertensive mice despite increased BBB permeability. Although we could not associate the protective effect offered by the microglia depletion with an improvement of myelin intensity in sites of BBB leakages, we occasionally observed impaired neuronal tracts (Thy1^+^) at the site of leakages, which may account for the decline in short-term memory induced by Angiotensin II. The high variability of the locations and sizes of the BBB leakages is however not compatible with neuronal tracing of Thy1^+^ neurons. Furthermore, the observed beneficial effect of PLX5622 might also result from a preserved neuronal plasticity as a recent study has revealed that the downregulation of neuronal and synaptic genes in an Alzheimer’disease model was prevented in absence of microglia [47].

Microglia depletion can be achieved by genetic deletion (*Csf1r^-/-^ Pu. 1^-/-^* mice), or by a pharmacologic depletion (CSF1R signaling inhibition) [29]. In the present study, we opted for pharmacological depletion as it is time-controlled and applicable in any mouse strain, using the highly selective CSF1R inhibitor PLX5622 for the depletion of microglia cells. PLX5622 has proven to induce a rapid and effective microglia depletion of more than 90% after 3-7 days of treatment [41, 42, 48, 49]. Even after 3 months of PLX5622 treatment cortical Iba-1^+^Cx3Cr1^+^ cell numbers were reduced by 90% (Fig. 2B), a finding that was confirmed on whole brain by flow cytometry of CD45^int^Cx3Cr1^hi^CD11b^hi^ cells. These data are in line with previous findings using CSF1R inhibition in several mouse models [41, 50]. While tissue macrophages and circulating monocytes express CSF1R as well, their survival is not only depending on CSF1lCSF1R signalling but relies also on CCL2/CCR2 signalling which is not present in microglia. We expected therefore that the PLX5622 treatment would lead to a mild reduction of tissue macrophages and monocyte populations [51]. Perivascular macrophages (PVMs) are brain macrophages that originate from hematopoietic precursors cells, express the mannose receptor (CD206^+^) and reside in the perivascular space of penetrating vessels [52, 53]. Activation of PVMs subsequent to BBB leakage has been shown to induce the production of ROS [38, 54]. Their depletion, using the injection of clodronate liposomes, was able to preserve both short-and long-term memory in a spontaneous hypertensive mouse model [38]. Although there was no change in PVM numbers following Ang II infusion in our study, as observed in earlier work [38] we found however a 50% reduction of the CD206^+^ PVM numbers due to the PLX5622 treatment. We cannot exclude that the favorable cognitive outcome in the hypertensive group after PLX5622 treatment is partly due to a reduction in the number of PVMs, although the total number of PVMs, limited to the perivascular space of large vessels, is clearly inferior to the > 2 million microglia populating a mouse brain [55]. To further investigate the effect of long-term CSF1R inhibition on circulating immune cells, we performed flow cytometry of the peripheral blood. We found a reduction of more than 50% of the non-classical Ly6C^low^ monocytes in the PLX5622 treated groups while there was no effect of the treatment on the classical Ly6C^high^ monocytes. Comparable results were obtained in a mouse study using PLX5622 for 7-days with a 30% reduction of the non-classical monocytes (Ly6C^Low^) and no effects on the classical monocytes (Ly6C^Hi^)[48]. In another study using a CSF1R antibody in cynomolgus monkeys, a decrease in non-classical CD14^+^CD16^+^ monocytes was also observed while the classical CD14^+^CD16^−^ monocyte population was unchanged [56]. Overall, the selective partial depletion of non-classical monocytes is in line with their low/negative CCR2 expression in comparison to classical monocytes (CCR2^high^), making them vulnerable to CSF1R inhibition. There is in the literature no evidence that non-classical monocytes are directly involved in the pathogenesis of cerebrovascular disease or that depletion of the non-classical monocytes is protective in this disease.

In addition, the depletion of microglia, and partially PVMs as achieved in our study, did not lead to a blood pressure reduction. The contribution of both macrophage subsets in the modulation of the autonomic activity in cardiovascular regulatory centers is well known [35, 57-61]. Previous studies reported that Ang II was able to increase blood pressure via the activation of microglia due to a lower BBB permeability in the paraventricular nucleus of the hypothalamus (PVN) in hypertensive animals [57, 62]. We did not observe a decrease of BP after microglia depletion, possibly because microglia were already depleted prior to, not after [62], Ang II-induced neurogenic hypertension.

The use of the *Cx3Cr1*^GFP/WT^ x *Thy1*^YFP/0^ mouse model may have lead to a reduction in Cx3Cr1 levels. As the insertion of GFP in the Cx3cr1^GFP^ mouse model was performed at the expense of 390 base pairs in the second exon of the Cx3Cr1 gene, its interaction with its ligand Cx3CL1 is altered as demonstrated when the mouse line was developed [64]. As a result, only heterozygous mice - as in the present study - should be studied to understand the behavior of microglia under healthy and pathological conditions [63].

In summary, we have shown that short-term memory impairment induced by prolonged Ang II infusion is absent when microglia are depleted using a CSF1R inhibitor. Cognitive effects of PLX5622 treatment were independent of changes in cardiovascular function and blood brain barrier permeability. This novel finding supports the hypothesis that microglia play a critical role in the pathogenesis of hypertension related cognitive impairment. This is a major step towards the development of theranostics targeting the CSF1R (e.g. [^11^C]CPPC[65]). An adequate modulation of microglia density and phenotype may constitute a relevant approach to prevent and/or limit the progression of vascular cognitive impairment.

## Acknowledgments

Plexxikon unrestrictedly contributed to the study by supplying the control and PLX5622 diets. We are grateful to Lieve Temmerman and Lou Maas for their technical assistance with the genotyping of the mouse model.

## Declarations of interest

None

## Funding information

This project has received funding from the European Union’s Horizon 2020 research and innovation programme under grant agreement No 666881, SVDs@target and from a Research Grant of the European Society of Hypertension and Servier attributed to S. Foulquier.

## Author contribution statement

DK, TU, JP, EAB, RJvO and SF contributed to the conception and design of the study. DK, BTvH, IVM, KJS, HvE, EW, PG and SF performed acquisition and data analysis. DK, PG, WMB, JP, EAB, RJvO and SF wrote the manuscript.

## Abbreviations list

Ang II: Angiotensin II
BBB: blood-brain barrier
CSF1: colony stimulating factor 1
CSF1R: colony stimulating factor 1 receptor
cSVD: cerebral Small Vessel Disease
MBP: myelin basic protein
MRI: magnetic resonance imaging
MWM: Morris water maze
OLT: object location task
PVM: perivascular macrophages
SBP: systolic blood pressure
SEM: standard error of the mean
TBS: tris buffered saline
VCI: vascular cognitive impairment
WMH: white matter hyperintensities.

